# ratioPCR delivers precise real-time quantification from multi-template PCR via mechanistic bias correction

**DOI:** 10.1101/2025.11.20.686216

**Authors:** Jochen Kinter, Adeline Stiefvater, Samira Osterop, Michael Sinnreich

## Abstract

Quantitative PCR (qPCR) is widely used for nucleic acid quantification but often lacks precision in detecting subtle relative changes, whereas digital PCR offers higher accuracy at reduced throughput and accessibility. Here, we introduce ratioPCR (rPCR), a bias-corrected qPCR approach that quantifies target ratios in real time by analyzing the fluorescence signal ratios during coupled amplification. Employing a mechanistic calibration model, rPCR corrects for differences in amplification efficiency, fluorophore intensity, and probe cross-reactivity, thereby achieving near-theoretical accuracy on standard qPCR platforms. Applied to alternative splicing analysis in a myotonic dystrophy type 1 model, rPCR accurately quantified isoform ratios and detected drug-induced splicing rescues in a high-throughput format. This method offers a fast, cost-effective, and scalable solution for precise relative nucleic acid quantification with potential applications in genotyping, mutation frequency estimation, and copy number variation analysis.

## INTRODUCTION

Quantitative PCR (qPCR) is a foundational tool in molecular biology that enables rapid, sensitive, and specific nucleic acid quantification in research, diagnostics, and environmental monitoring^1,2^. Despite its versatility, qPCR accuracy is strongly influenced by RNA quality, assay design, reference standards, and reaction conditions, which limit reproducibility^3,4^. Digital PCR (dPCR) addresses some of these challenges by providing absolute quantification without calibration curves and a greater tolerance to inhibitors^5–7^. However, dPCR has a higher cost, lower throughput, and a reduced dynamic range, constraining its use^8^. For many applications, qPCR remains the method of choice; however, it struggles to reliably detect subtle relative changes, often below twofold, reliably^9^.

Shortly after the invention of PCR, competitive PCR was introduced to enable relative quantification by co-amplifying two templates with a shared primer pair^10–13^. Competitive PCR belongs to the class of multi-template PCR, which is characterized by the co-amplification of two or more targets with a common primer pair, in contrast to multiplex PCR, in which each target is amplified using its own primer set. Multi-template PCR has been applied in several fields, including environmental microbiology for analyzing complex microbial communities^14^, molecular evolution^15^, general library preparation, DNA storage^16^, mutation analysis, and CRISPR-Cas9 editing efficiency^17^.

Although conceptually powerful, competitive PCR suffers from amplification biases caused by differences in amplification efficiency and artifact formation, such as heteroduplexes and chimeras. These effects can distort the measured ratios and limit the accurate quantification^18,19^. Although dual-probe competitive PCR and empirical bias corrections have been reported^20–22^, a mechanistic correction framework addressing probe imbalance, cross-reactivity, and ratio-dependent drift is lacking. Without addressing this issue, a method combining high-throughput, low-cost, and accurate quantification of small relative differences remains lacking, representing a bottleneck for biomarker validation, clinical testing, and drug discovery.

To address this, we developed ratioPCR (rPCR), which determines the relative target abundance directly from real-time fluorescence signal ratios during coupled amplification, avoiding reliance on error-prone cycle threshold (Cq) values. We established a detailed kinetic model to understand the mechanisms leading to the bias observed in the multi-template PCR. Based on this knowledge, we set up a simplified analytical calibration model that corrects for amplification efficiency differences, fluorophore intensity disparities, and probe hybridization biases, thereby enabling accurate and reproducible ratio measurements on standard qPCR platforms.

As a proof of concept, we applied rPCR to quantify alternative splicing in myotonic dystrophy type 1 (DM1), a disease in which sequestration of the splicing factor MBNL1 by expanded CTG repeats leads to widespread mis-splicing^23,24^. Accurate high-throughput quantification of splicing biomarkers is essential for mechanistic studies and therapeutic screening. We demonstrated that rPCR quantifies isoform ratios and detects drug-induced splicing rescue in a DM1 cellular model.

Together, these results establish rPCR as a robust, cost-effective, and scalable method for relative nucleic acid quantification, bridging a methodological gap between qPCR and digital PCR and opening new avenues for splicing analysis, biomarker quantification, and therapeutic screening.

## MATERIAL AND METHODS

### Primer and probe design

We selected three genes, KIF13A, *NUMA1*, and *SYNE1*, which are known to be misspliced in myotonic dystrophy type 1 (DM1), for method development^25–27^ (Supplementary Figure 2A). Using RNA from DM1 patient-derived and control myoblasts, RT-PCR confirmed the relative exon exclusion in DM1 (Supplementary Figure 2A). Agarose gels revealed additional slower-migrating bands corresponding to heteroduplexes between isoforms (Supplementary Figure 2B and Supplementary Figure 3), which complicate gel-based quantification but do not affect probe-based detection.

Inclusion-specific probes matched the alternatively spliced exon, whereas exclusion-specific probes spanned the exon–intron junction, allowing partial hybridization to the inclusion isoform. Gibbs free energy calculations predicted higher cross-reactivity for the exclusion probes than for the inclusion probes (Supplementary Figure 4A). Probe specificity testing with synthetic isoform oligonucleotides confirmed these predictions: inclusion probes showed no detectable cross-reactivity, whereas exclusion probes exhibited variable cross-reactivity, most pronounced for *NUMA1* (Supplementary Figure 4B). The thermodynamic predictions (ΔG) closely matched the experimental results, supporting their use in the probe design.

### Oligonucleotides

Primers, probes and template oligonucleotides were purchased from Integrated DNA Technologies and Microsynth. For details, see Supplementary Table. For calibration experiments, either long single-stranded oligonucleotides (Ultramer DNA oligos, IDTDNA) or double-stranded DNA fragments (gBlock Gene Fragments, IDTDNA) were used. Reference samples were obtained by mixing the defined ratios of the two targets of interest.

### rPCR

All PCR reactions were performed with a Quantstudio 5 PCR machine from ThermoFisher equipped with a 384-well plate-compatible heating block. The Luna Universal One-Step RT-qPCR system (New England Biolabs) was used to perform reverse transcription and amplification in a single step. Reverse transcription was omitted from experiments using synthetic reference samples. The final primer and probe concentrations were 0.4 µM primer and 0.4 µM probes for *NUMA1* and KIF13A assays, 0.2 µM primer and 0.2 µM probes for the *SYNE1* assay, if not stated differentially. The cycling protocol was as follows: 10 min at 55 °C and 10 min at 95 °C, followed by 45 cycles of 10 s at 95 °C, and 45 s at 60 °C. The measured fluorescence values were exported as an Excel file and imported into R software for further analysis.

### dPCR

dPCR was conducted using the QIAcuity One, 5plex platform system (QIAGEN). The reaction was run in QIAcuity Nanoplates 8.5k 96-well plates (QIAGEN). The cycling conditions were 95□°C for 10□min, followed by 45 cycles of 95□°C for 15□s and 60□°C for 54□s.

### Calibration and Evaluation of the Analysis Method

An R package was developed to enable fast and simple analysis of ratioPCR data. To obtain the calibration parameters, the experimental data for the reference samples were fitted to Eq. Two using the nlsLM function from the R package minipack.lm (see Supplementary Notes).

This calibration was performed once per assay, and the resulting parameters were then applied to the measured apparent ratio in Eq. 2 to calculate the true ratio.

### Kinetic Modeling of rPCR

For the development of a kinetic model of rPCR OpenModelica^28,29^ 4.0, the Biochem^30^ 2.0 library was used. Postprocessing and visualization of the simulation results were performed in R (see Supplementary Notes).

### Cell culture

Immortalized human myoblasts were obtained from the Institute of Myology, Paris,^31^ and were cultured in Growth Media (40% Skeletal Muscle Cell Growth Medium Kit (C-23160-PRO PromoCell), 40% Nutrient Mixture F-10 Ham medium (Sigma N6013), and 20% FBS (Gibco 10270 106) supplemented with 50 µg/mL gentamicin (Sigma G1272-100ML). For differentiation, the cells were plated in a 96-well plate at a density of 25’000 cells per well for two days in proliferation media (Skeletal Muscle Cell Differentiation Medium (C-23061-PRO PromoCell and 50 µg/mL gentamicin). After an additional 3 days in differentiation medium, the cells were harvested and RNA was purified.

### RNA purification

After washing the cells with PBS, 150 µL of lysis buffer (4 M GTC, 50 mM Tris-HCl pH 7.5, and 25 mM EDTA) was added to each well for 5 min. A total of 150 µL of 70% ethanol was added per well and mixed. The solutions were transferred to a 96 well silica based nucleic acid purification plate (Axygen, VWR). After removing the solution using a vacuum manifold the wells were washed twice with a GTC wash buffer (1 M GTC, 12.5 mM Tris-HCl pH 7.5, 6.25 mM EDTA) and twice with a RNA wash buffer (60 mM potassium acetate, 10 mM Tris-HCl pH 7.5, 60% EtOH). After centrifugation of the plate at 4000 × g to remove residual liquid, RNA was eluted using 50 µL water.

## RESULTS

### rPCR concept and assay design

We developed ratioPCR (rPCR), a quantitative PCR approach that enables the rapid and accurate measurement of relative nucleic acid abundance. rPCR co-amplified multiple targets using a shared primer pair, generating a coupled amplification (Figure 1A). Template-specific hydrolysis probes detect each target, and instead of relying on the cycle threshold (Cq), rPCR monitors the ratio of fluorescence signals in real-time (Figure 1B,C). The template ratio was captured by the real-time ratio of isoform-specific probe signals and accurately corrected using a calibration model (Figure 1C, Supplementary Figure 1).

**Figure 1:**
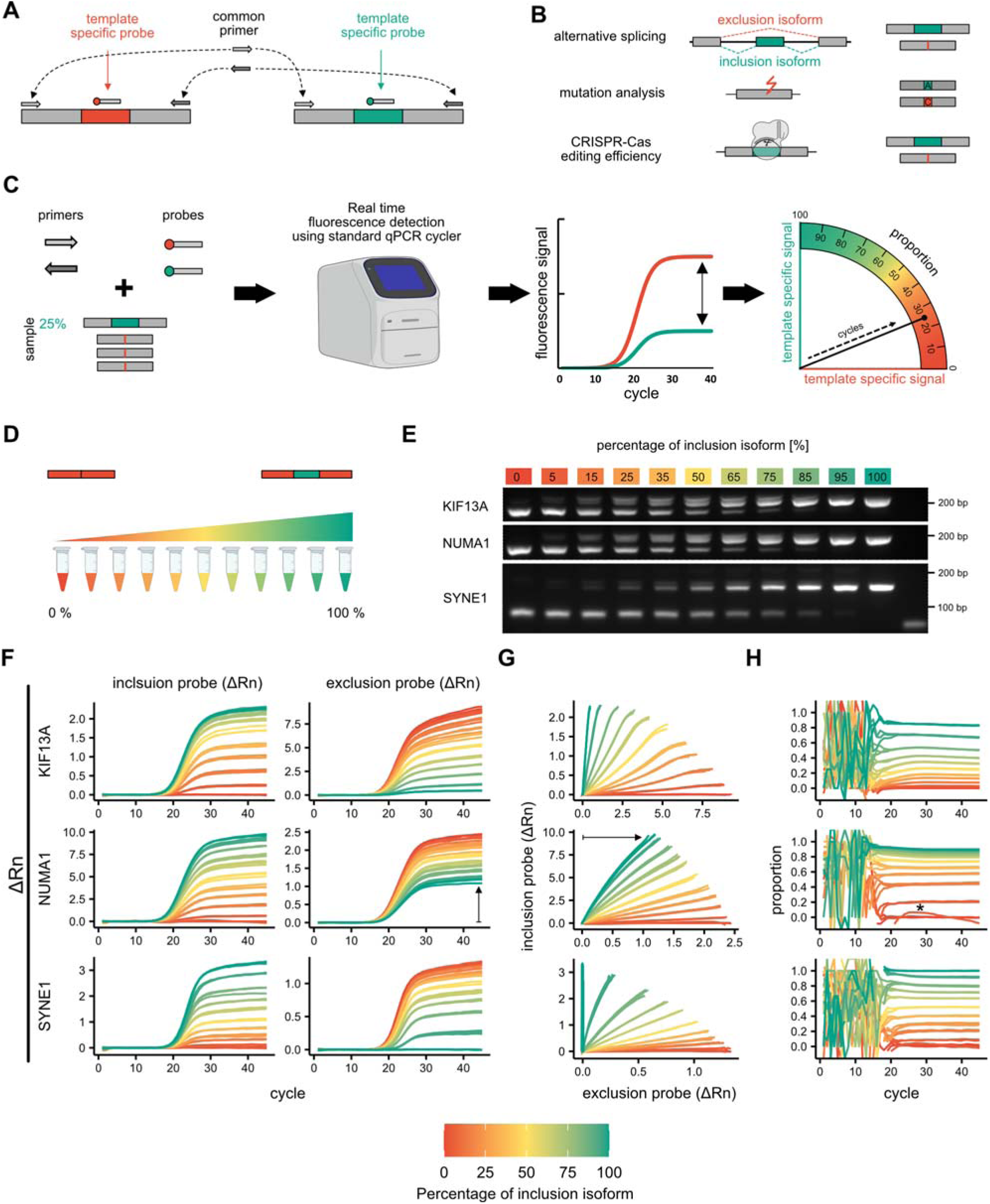
Principle of ratioPCR (rPCR) and its validation using reference samples demonstrates assay specific biases. **a**, Alternative splicing generates distinct mRNA isoforms containing different exon combinations. **b**, rPCR uses a common primer pair to co-amplify both isoforms, coupled with two isoform-specific hydrolysis probes labeled with distinct fluorophores for inclusion and exclusion detection. **c**, During rPCR, amplification is monitored via fluorescence. The ratio of probe signals reflects the biased isoform proportions and is quantified using a standard qPCR instrument. **d**, Genomic context and assay design for three alternatively spliced genes (KIF13A, *NUMA1*, *SYNE1*). Red boxes mark the alternative exons. Primer binding sites (arrows) and probe locations (red: inclusion; green: exclusion) are shown. **e**, Agarose gel electrophoresis of RT–PCR products from control and DM1 myoblast RNA reveals altered splicing patterns. Heteroduplexes between isoforms form slower-migrating bands. **f**, Defined mixtures of synthetic inclusion and exclusion oligonucleotides were prepared to generate reference samples with known isoform ratios. **g**, RT–PCR of the reference samples followed by agarose gel electrophoresis shows isoform-specific banding patterns for KIF13A, *NUMA1*, and *SYNE1*, confirming the expected inclusion/exclusion shifts. **c-h**, Real-time fluorescence signals were analyzed to assess ratio dynamics and assay performance. Fluorescence curves are shown as raw Rn signals (**c-e**) and baseline-subtracted ΔRn signals (**f-h**). Data visualizations include: classical amplification plots (**c,f**), slope plots displaying the fluorescence of both probes against each other (**d,g**), ratio plots illustrating the evolution of the measured isoform proportion during amplification (**e,h**). All reactions were run in technical triplicates. A clear signal distortion due to cross-reactivity of the *NUMA1* exclusion probe is indicated by an arrow. A background subtraction artifact leading to severe ratio distortion in one reaction is marked with an asterisk.

To illustrate feasibility, we applied rPCR to quantify alternative splicing using primers flanking an alternatively spliced exon to co-amplify both isoforms, while isoform-specific probes distinguished inclusion and exclusion variants (Figure 2B,C). We designed assays for three genes (KIF13A, *NUMA1*, and *SYNE1*) known to be mis-spliced in myotonic dystrophy type 1 (DM1)^25–27^ (Supplementary Figure 2A). RT–PCR of RNA from DM1 and control myoblasts showed differential splicing for all three genes (Supplementary Figure 2B) and revealed heteroduplexes, which appeared as slower-migrating bands due to rehybridization of isoforms (Supplementary Figure 2B & 3).

**Figure 2:**
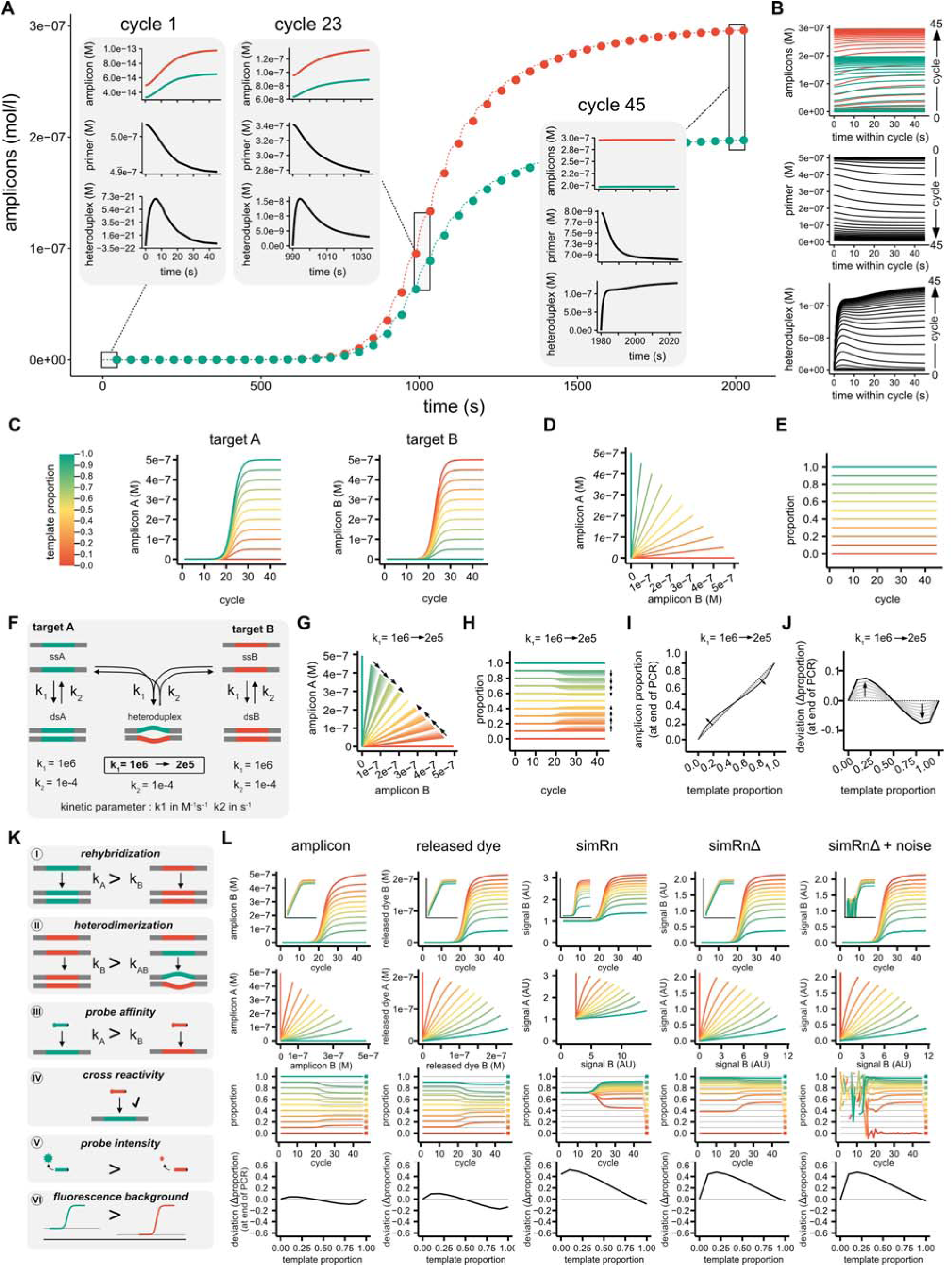
Simulation using the kinetic rPCR model. **a**, Example of a kinetic rPCR simulation with an initial template proportion of 40% target A (green) and 60% target B (red), starting from 1×10□ total copies. Amplicon concentrations for targets A and B are shown over time. Dots indicate concentrations at the end of each PCR cycle. Insets show detailed profiles of amplicons, primers, and heteroduplexes at cycles 1, 23, and 45. **b**, Time-course profiles of amplicons, primers, and heteroduplexes within all PCR cycles, illustrating how molecular dynamics evolve throughout the reaction. **c-e**, Simulated run under idealized conditions with equal rehybridization and heterodimer formation rates, and no probes. Amplicon concentrations of both targets maintain a linear relationship (**d**). No template-to-product bias is observed under these conditions (**e**). **f-j**, Simulations highlighting the impact of heteroduplex formation on ratio distortion. Schematic of rehybridization and heterodimerization kinetics (**f**). **g**, The linear relationship between targets is disrupted for input ratios other than 0, 0.5, or 1. h, Progressive shift of the ratio toward 1:1 is observed during amplification. **i**, Comparison of initial template ratio and final amplicon ratio shows systematic bias dependent on heterodimer kinetics. **j**, The magnitude and direction of ratio distortion depends on the starting proportion. **k-l**, Simulations incorporating multiple sources of rPCR bias and measurement distortion. **k**, Illustration of factors affecting rPCR accuracy: (**I**) Different rehybridization rates → asymmetric amplification efficiency, (**II**) Heteroduplex kinetics → convergence toward equimolar ratios, (**III**) Probe affinity differences → amplification and signal drift, (**IV**) Probe cross-reactivity → target misassignment, (**V**) Fluorescence intensity differences and instrument sensitivity → signal distortion, (**VI**) Background levels and noise impact the finally measured fluorescence signals. **l**, Multilevel output simulation showing: amplicon concentration, released dye concentration, fluorescence readout (Rn), background-subtracted signal (ΔRn) without and with noise. Each level introduces increasing deviation from the original template ratio which is demonstrated by amplification graphs of target b (1st row), slope graphs illustration non-linear relationship between target A and B (2nd row), ratio plot illustrating different ratio bias pattern (3rd row), and deviation at the end of the PCR from the starting template ratios (4^th^ row).

We evaluated probe performance both computationally and experimentally. The predicted Gibbs free energy (Supplementary Figure 4A) suggests high specificity for the inclusion probes and variable cross-reactivity for the exclusion probes. Experimental testing using synthetic templates validated these predictions: inclusion probes showed no off-target signal, whereas exclusion probes, particularly for *NUMA1*, displayed notable cross-reactivity (Supplementary Figure 4B).

### Validation Using Reference Standards

To evaluate analytical accuracy, synthetic reference samples were prepared by mixing known ratios of inclusion and exclusion templates (Figure 1D). The amplification produced the expected isoform patterns, as visualized by gel electrophoresis (Figure 1E). Real-time fluorescence signals were recorded as normalized fluorescence (Rn) and baseline-subtracted (ΔRn) signals. These data were visualized using amplification curves, slope plots (inclusion vs. exclusion signal), and proportion plots tracking the proportion of inclusion signals over the cycles (Figure 1F-H).

The apparent proportion (ψ_a_) of inclusions to the total signal, also referred to as the percent/proportion spliced in (PSI or ψ), was calculated as follows:

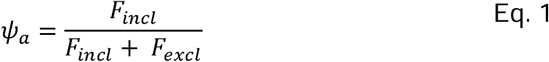

In rPCR, higher inclusion template levels resulted in increased inclusion and decreased exclusion signals (Figure 1F). Slope plots revealed a non-linear relationship between signals, indicating a bias in probe hydrolysis and/or amplification efficiency. The proportion plots showed dynamic drift in the apparent proportion (ψ□), highlighting systematic deviations. For *NUMA1*, the cross-reactivity of the exclusion probe was evident even in samples containing only inclusion templates (arrow, Figure 1f,g). Additionally, one replicate displayed an erroneous ΔRn-based ratio owing to incorrect automatic background subtraction (asterisk, Figure 1H). These findings underscore the need for calibration to correct systematic distortions.

### Kinetic Modeling Reveals Sources of Bias

To mechanistically dissect amplification bias in rPCR, we developed a detailed kinetic model using OpenModelica^28,32^ and BioChem library^30^. This model simulates real-time PCR as a system of biochemical reactions, including primer-template hybridization, strand synthesis, rehybridization, heteroduplex formation, probe hybridization, and fluorescence generation (Figure 2, Supplementary Notes). The system comprises over 250 differential equations with kinetic starting parameters sourced from published data^33,34^.

Under idealized simulation conditions, equal rehybridization and heteroduplex formation rates and no probes and template ratios were preserved across all cycles (Figure 2C-E). In contrast, introducing unequal rehybridization kinetics led to progressive distortion, with ratios drifting toward 1:1 (Figure 2F-J). These simulations recapitulate known phenomena in multi-template PCR^10,18,35^ and validate the predictive capacity of the model. Moreover, the model predicted that mixed-template reactions can exhibit higher apparent efficiency than single-template reactions owing to heteroduplex dynamics (Supplementary Figure 5).

Simulations also explored how factors such as probe cross-reactivity, fluorophore intensity differences, and background noise distorted the observed signal ratios (Figure 2K-L). These findings motivated the development of a simplified analytical model to support calibration and correction.

### Analytical Model for Calibration

We derived an analytical model that maps the true proportion ψ to the apparent ψ□ via a small set of interpretable parameters: α (relative probe intensity/efficiency), r□/r□ (inclusion/exclusion probe cross-reactivity), and γ (ratio-dependent amplification term capturing rehybridization driven ratio-drift). The compact form used throughout is:

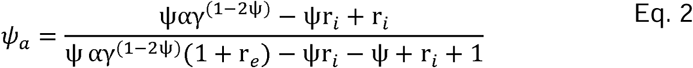

with variable definitions in the caption and derivations in Supplementary Note 1.

The sensitivity analysis showed that modest deviations in any parameter could produce substantial nonlinear distortions (Figure 3A). The model was fitted to endpoint fluorescence values from the reference samples using nonlinear regression. Assay-specific calibration parameters were estimated from the Rn and ΔRn data. The corrected values (ψIZl) closely matched the theoretical ratios (ψ_t_), substantially reducing the residual error (Figure 3B). The model robustly captured key bias mechanisms and improved the assay precision.

**Figure 3:**
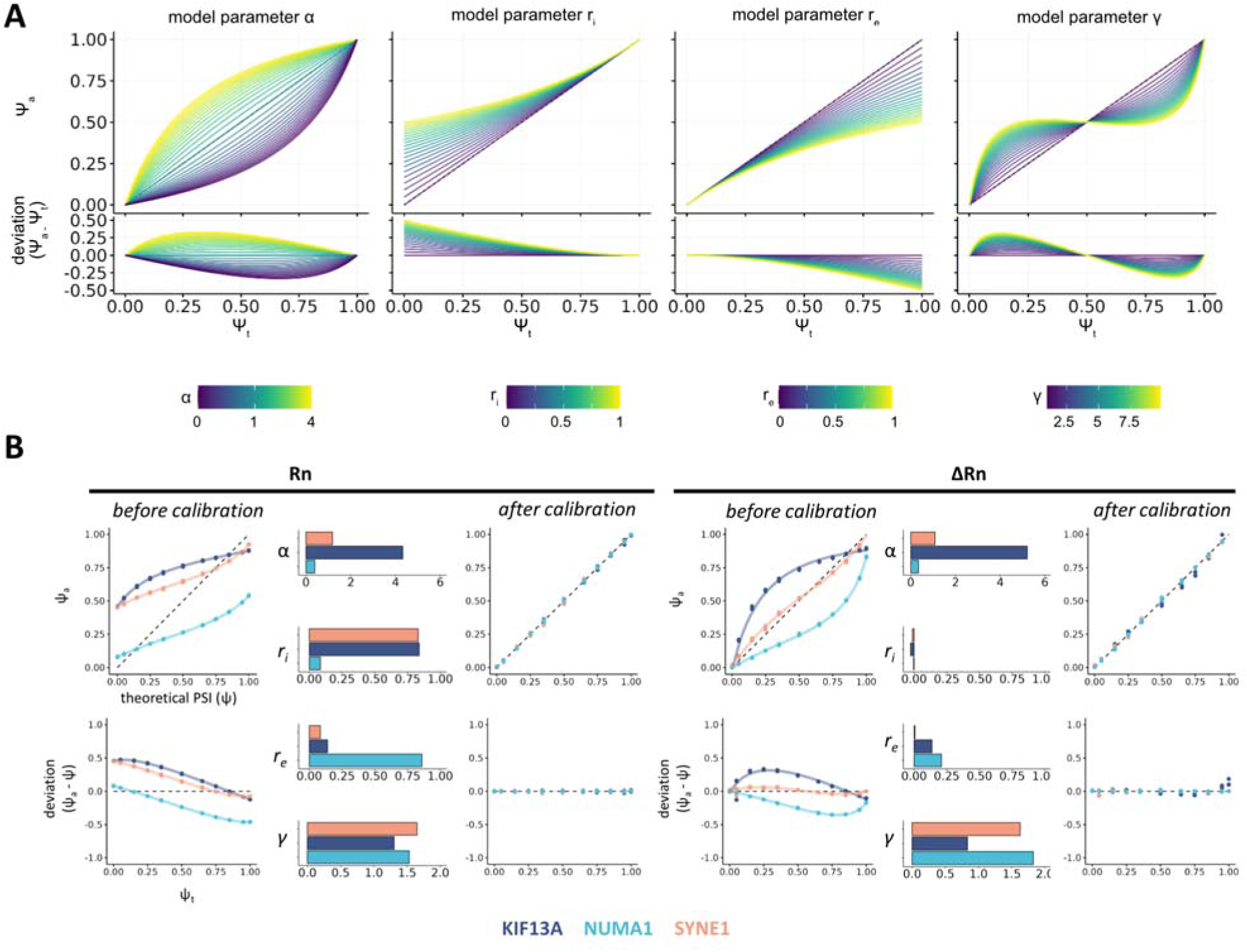
Calibration of rPCR using the analytical bias correction model. **a,** Sensitivity analysis illustrating how variation in model parameters (α, γ, r□, r□) influences the apparent isoform ratio (ψ□) relative to the true input ratio (ψ□), based on simulations using the analytical bias model (Eq.□2). Each parameter reflects a distinct source of assay bias: α captures differential fluorescence intensities between probes; γ accounts for ratio-dependent amplification bias; r□ and r□ represent cross-reactivity of the exclusion and inclusion probes, respectively. ***b***, Experimental data from synthetic reference samples were fitted to Eq.□2 to extract assay-specific calibration parameters. The apparent ratios (ψ□) calculated from raw (Rn) and baseline-subtracted (ΔRn) fluorescence data were corrected using the model to yield calibrated ratios (ψ_c_). Residual plots show improved concordance with theoretical values after calibration, demonstrating effective bias correction (n=3).

#### Synchronizing amplifications identifies a reliable single-cycle window

Next, we evaluated whether individual PCR cycles beyond the final cycle could yield accurate isoform ratios. Because reactions reached maximal amplification at different cycles, we synchronized them using the cycle with maximal fluorescence increase (Ci), defined as cycle 0 (Figure 4a). We analyzed the relative cycles (C_r_) from −10 to +15 (Figure 4A), comparing the calibrated and uncalibrated ψ values using Rn, ΔRn, and first-derivative signals (derRn) (Figure 4B). Calibration improved the accuracy across a wide cycle range (Figure 4B), with derivative-based signals yielding optimal results near Ci, whereas Rn and ΔRn signals performed best at later cycles (Figure 4C,D).

**Figure 4:**
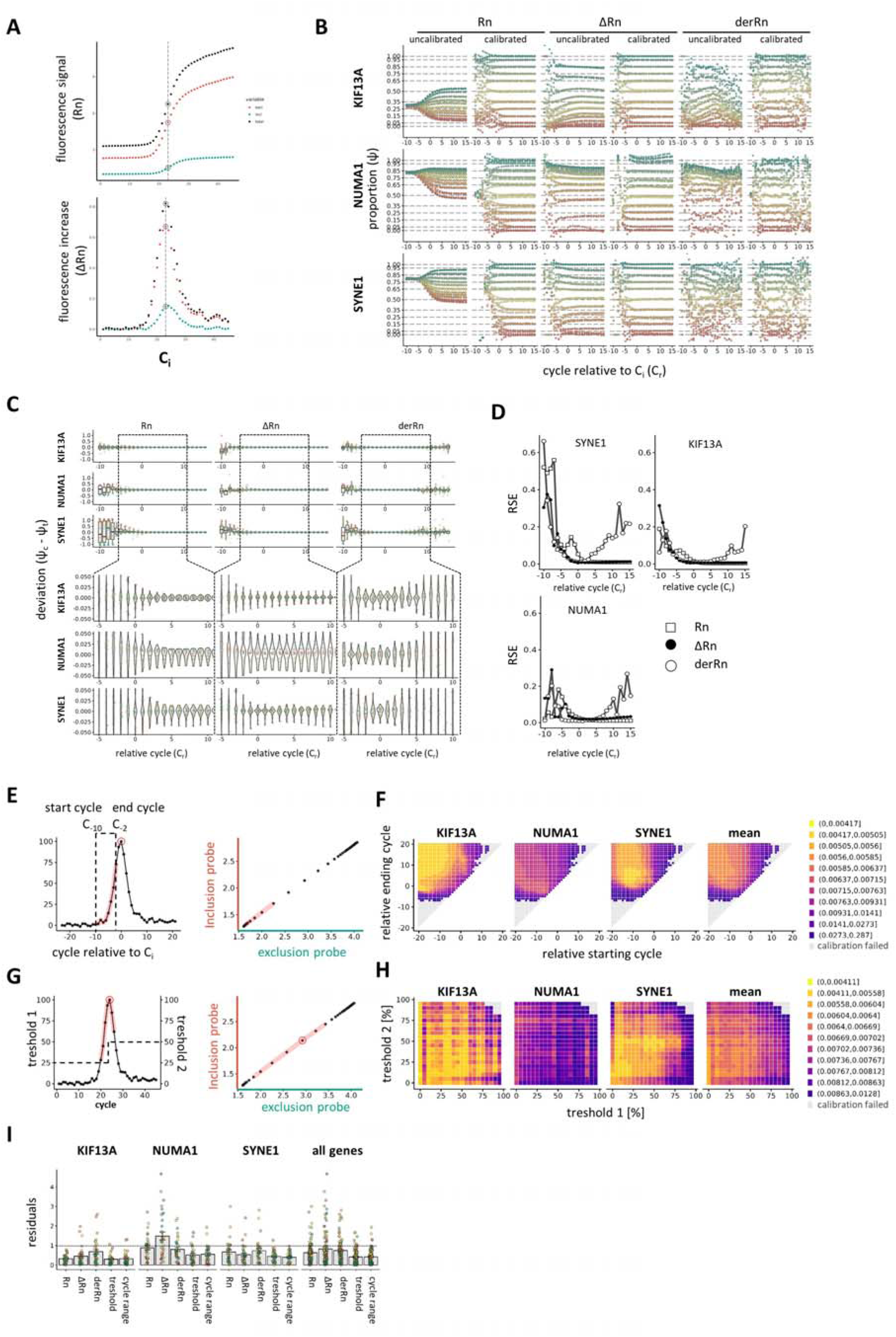
Ratio quantification by slope calculation using a defined range of cycles. **a**, Example of cycle alignment using the derivative of the Rn fluorescence signal. The cycle with the maximum fluorescence increase (C_i_) is defined as cycle 0. **b**, Apparent isoform proportions (ψ_a_) and calibrated proportions (ψ_c_) were calculated for each cycle from C□□□ to C□□ relative to C□. Three signal types were evaluated: raw fluorescence (Rn), baseline-subtracted signal (ΔRn), and the first derivative of Rn (derRn). Horizontal dashed lines indicate theoretical isoform proportions. **c**, Residuals between calibrated proportions and theoretical values at each cycle demonstrate the effect of cycle position and signal type on calibration accuracy. **d**, Relative standard error (RSE) of calibrated ratios across cycles. Lower RSE indicates better precision. **e-h** Two different approaches to calculate a ratio over a range of cycles were applied. **e**,**f,** The range was defined by a starting and ending cycle relative to the maximum of the first derivate of the signal. **f**, Matrices showing the relative standard error for ranges with different relative start and end cycles. Matrices are shown for KIF13A, *NUMA1*, *SYNE1*, and the average of all three assays. **g**,**h**, Threshold method defines the cycles by setting a threshold for the first derivative of the Rn signal (derRn). Only cycles with derRn signals above the thresholds before and after the maximum are used for the slope calculation. **h**, Matrices visualizing the relative standard error for different threshold combinations. Matrices are shown for *KIF13A*, *NUMA1*, *SYNE1*, and the average of all three assays. **i**, Comparison of the absolute residuals of different methods to calculate the apparent ratio. Three methods using only one defined cycle for calculation (Rn, ΔRn, and derRn) was compared to methods based on slope calculation in a range of cycles using the threshold method or defined cycle range. All reactions were run in technical triplicates.

#### Multi-cycle slope fitting provides modest but consistent gains

To further improve the accuracy, we tested the multicycle slope fitting across defined cycle ranges. Two strategies were employed: (1) relative positioning around Ci, and (2) thresholding based on the fluorescence derivative. For each cycle range, the inclusion/exclusion slope was calculated and used to estimate ψ□, which was then calibrated (Figure 4E,G).

The accuracy maps revealed the optimal cycle range for each assay (Figure 4F,H). A final comparison of all strategies using the residual standard error as an accuracy indicator showed that multicycle slope fitting offered slightly improved accuracy and robustness over single-cycle methods, particularly when using Rn signals (Figure 4I). Across all assays, the mean deviation from the theoretical ratios was <1%.

#### rPCR approaches dPCR precision while preserving qPCR throughput

To benchmark the rPCR against an established standard, we compared it with digital PCR (dPCR) using synthetic reference samples. For *NUMA1*, dPCR 1D/2D partition plots showed the expected two well-separated populations in single-isoform samples and a prominent intermediate population in 1:1 mixes (appearing as additional “rain” but here reflecting mixed-template partitions, Figure 5A asterisk). Normalized fluorescence confirmed a higher total signal and intermediate inclusion proportion in mixed partitions (Figure 5B,C), consistent with kinetic model predictions (Supplementary Figure 5), and density profiles resolved discrete mixed subpopulations (≈1:1, 2:1, 1:2; Figure 5D).

**Figure 5:**
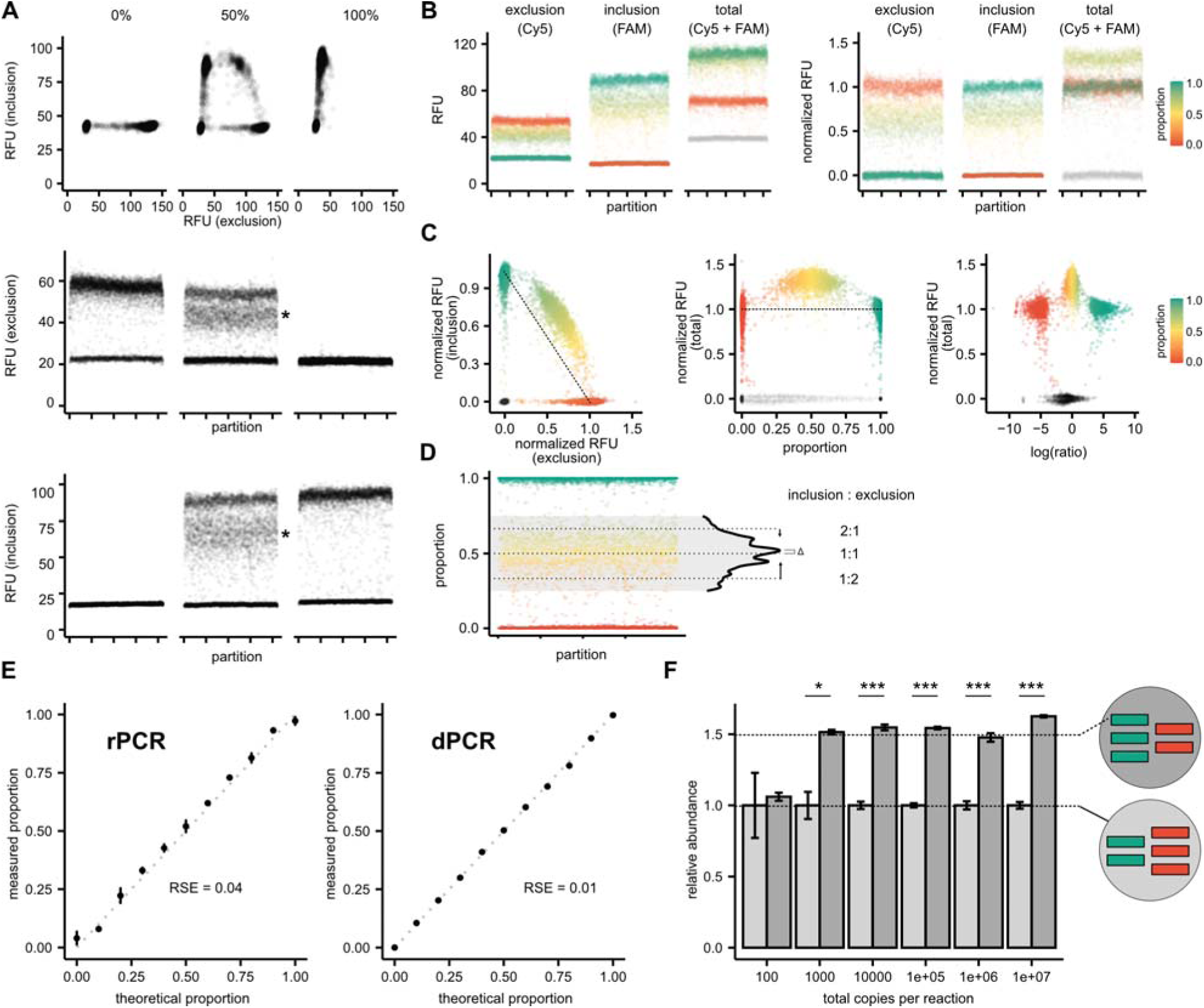
Comparison of dPCR and rPCR. **a**, 1D and 2D plots of dPCR experiments using *NUMA1* reference samples with only exclusion (0%), only inclusion isoform (100%), and a 1:1 isoform mixture. In the 1D plots both single isoform samples mainly, two populations are detected and a few partitions with medium fluorescence levels known as “rain”. In the mixed sample an increase of medium level fluorescence population is detected in the 1D plots, reflecting partitions containing both isoforms (see asterisk). **b**, The relative fluorescence units (RFU, left panel) were shown for both channels and their sum. Normalized RFU values for each channel and their sum are shown in the right panel. The color code reflect the proportion of the inclusion isoform [proportion = (nRFUincl / (nRFUFincl + nRFUexcl)]. **c**, Different graphical presentation of the mixed sample in 2D plot, proportion vs the total normalized RFUs, and the log(ratio) vs the total normalized RFUs demonstrating the increased total fluorescence levels for mixed template partitions. Due to heteroduplex formation, the amplification efficiency is elevated in mix template partitions. **d**, 1D partition plot indicating the distribution of different proportions. Most partitions have only one isoform. A density profile demonstrating distinct mixed population representing a 1:1, 2:1 or 1:2 isoform ratios. **e**, Comparison of rPCR with dPCR using KIF13A reference samples with known isoforms proportion. Both methods were able to accurately measure the proportions. rPCR performed in traditional qPCR machine reaching accuracy close to dPCR (Residual standard error, RSE_rPCR_ = 0.04 and RSE_dPCR_ = 0.01). **f**, Two samples with 40% and 60% of inclusion isoform, reflecting a 1.5 fold difference, were measured with different total copy numbers per reaction to analyze the limit of detection and the dynamic range to detect small changes (Welch two sample t-test: * p < 0.05, *** p < 0.0001). Panel **a-d** showing results from a *NUMA1* assay and **e-f** are generated using a KIF13A assay. All reactions were run in 4 technical replicates.

With the KIF13A references, rPCR (on a standard qPCR instrument) achieved RSE = 0.04, close to RSE = 0.01 for dPCR (Figure 5e). Sensitivity tests with 40% versus 60% inclusion (1.5-fold) across varying input copies showed that rPCR reliably discriminated subtle fold changes (fc=1.5) over a broad dynamic range (Welch’s two-sample t-test: p<0.05; **p<0.0001) (Figure 5f). Thus, rPCR can achieve dPCR precision for relative quantification while maintaining qPCR-level throughput and cost.

### Application to Disease Models and Screening

To demonstrate biological relevance, we applied rPCR to a cellular disease model of Myotonic Dystrophy type I^31^. We established a screening workflow to examine drugs for their potential to improve splicing missplicing in DM1 (Figure 6A). The final rPCR reactions were performed in 384-well plates using 10-µl reactions in a one-step format, combining reverse transcription and subsequent rPCR. All three assays detected consistent missplicing in DM1 cells with and without calibration (Figure 6B). We evaluated rPCR for screening applications by treating DM1 myoblasts with furamidine, a diamidine compound known to partially restore normal splicing. A dose-dependent rescue was observed for KIF13A and *SYNE1* with minimal changes in *NUMA1* expression (Figure 6C). These results support rPCR’s utility of rPCR for biomarker validation and drug screening in high-throughput formats, using standard qPCR platforms.

**Figure 6:**
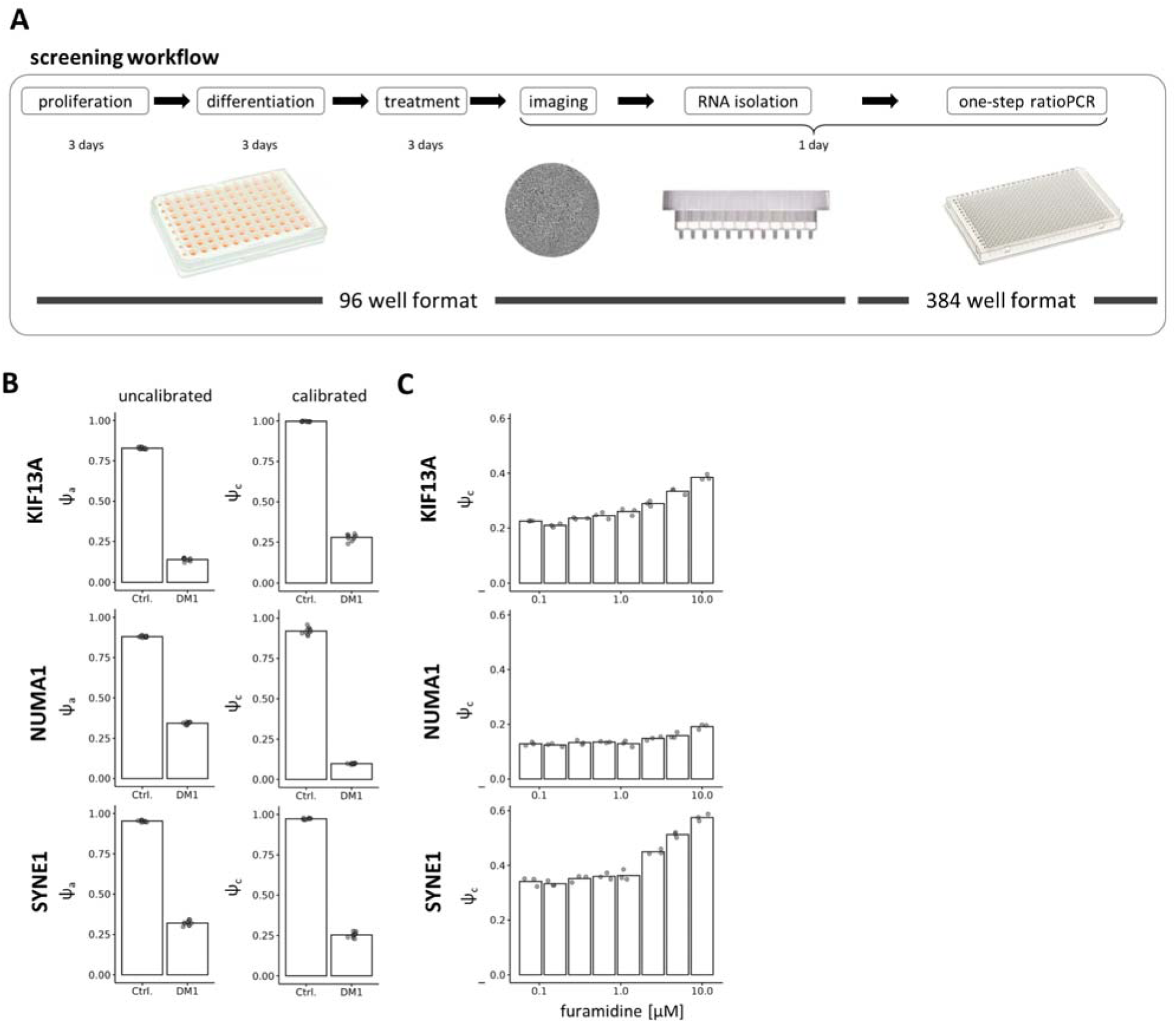
Application of rPCR to DM1 patient-derived myoblasts and drug response analysis. **a**, Workflow for the screening of compound effects on alternative mis-splicing in patient derived myotonic dystrophy type 1 differntitaed myotubes. Differentiated Myoblasts are imaged using a plate reader before RNA is isolated. RNA samples are directly used in a one-step rPCR. To enhance throughput rPCR is performed in a 384 format. **b**, Differentiated myoblasts from a myotonic dystrophy type 1 patient and a healthy control were analyzed using rPCR to quantify alternative splicing of three MBNL1-regulated transcripts (KIF13A, *NUMA1*, *SYNE1*). Uncalibrated (ψ□) and calibrated (ψ_c_) isoform proportions are shown for each gene. Both measurements detect pronounced splicing shifts in DM1 cells relative to controls, consistent with disease pathology. **c**, To assess rPCR performance in compound screening, DM1 cells were treated with increasing concentrations of furamidine, a diamidine compound previously shown to rescue mis-splicing in DM1. rPCR revealed a dose-dependent increase in exon inclusion for KIF13A and *SYNE1*, while *NUMA1* responded only weakly. Dots represent technical replicates (n≥3).

## DISCUSSION

rPCR addresses the pervasive gap between qPCR and dPCR: accurate, scalable, and cost-effective measurement of subtle relative differences in nucleic acid abundance. By reading out real-time fluorescence ratios under common primer amplification and applying mechanistic calibration that corrects probe- and amplification-related biases, rPCR achieves near-dPCR precision while preserving the throughput, economics, and accessibility of qPCR.

Our results show that during amplification, relative template ratios show nonlinear deviations in a ratio-and target-dependent manner, driven by multiple sources of bias. Two contributors dominate ratio distortion in multitemplate PCR: sequence-dependent rehybridization kinetics that differentially limit effective efficiency even with shared primers and heteroduplex formation that accelerates efficiency loss for abundant templates, yielding a well-described drift toward equimolarity^10,36^.

Our kinetic model reproduces these behaviors, including enhanced apparent efficiency in mixed-template contexts via heteroduplex dynamics and guided construction of a tractable analytical model with a handful of interpretable parameters (α, r□/r□, γ) suitable for routine fitting. The

We further optimized rPCR by systematically analyzing the signal types (Rn, ΔRn, and first derivative) and amplification cycles for ratio extraction. Our results show that synchronizing PCR curves using the cycle of maximum fluorescence increase (C_i_) enables a standardized comparison across samples. We demonstrated that single-cycle ratio estimates near C_i_, particularly using Rn signals, yield highly accurate results when calibrated. Moreover, analyzing the slope of isoform signal trajectories across multiple cycles slightly improves accuracy and robustness, with a consistent mean deviation from theoretical values of <1%.

The utility of rPCR was validated in a cellular model of myotonic dystrophy type 1 (DM1), in which mis-splicing of KIF13A, *NUMA1*, and *SYNE1* transcripts occurred. Application to furamidine-treated cells further demonstrated rPCR’s suitability of rPCR for compound screening, revealing dose-dependent correction of splicing defects. These findings suggest that rPCR is a versatile platform for both basic research and translational applications.

rPCR also clarifies a practical point regarding conventional qPCR: non-specific priming effectively converts a single-target assay into a multi-template reaction (a special case of competitive PCR when the specific and non-specific products co-amplify with the same primer pair). Our modeling and experiments indicate that this alters the apparent efficiency of a specific product by biasing both the curve shape and Cq. The bias scales with the fraction of non-specific products and accumulates over cycles—larger at low specific input and often negligible at high input. Crucially, even if nonspecific products are not detected by the probe, they compete for primers and distort the kinetics of the specific signal. Careful assay design to suppress non-specific amplification is essential.

Although rPCR can detect differences semi-quantitatively without calibration, assay-specific calibration with reference mixes is required to obtain accurate absolute ratios. This is practical because long sequence-verified oligonucleotides are widely available and affordable. The choice of fluorescence dye is also important, as a higher signal-to-noise ratio of the selected probe dyes will increase the assay resolution. Because rPCR has the advantage of detecting lower differences, it is difficult to accurately quantify very low/high template proportions (<1%, >99%). Our analytical model assumes shared primer-binding sites with comparable binding energetics; applications such as microbial or viral communities that use a common primer on heterogeneous priming sites may exhibit primer-site-driven bias that is not captured by the current parameterization.

A variety of methods exist for quantifying alternative splicing, including RT–PCR with gel electrophoresis□^37^, digital PCR^38,39^, MLPA^38^, or RNA-seq^40,41^. Although RNA-seq provides global insights, its high cost, data burden, and limited sensitivity for low-abundance isoforms make it impractical for large-scale or high-throughput applications. rPCR In contrast, rPCR enables the quantification of relative abundance using standard qPCR equipment with minimal computational overhead. Compared to digital PCR (dPCR), rPCR offers higher throughput and lower cost while maintaining high precision for relative quantification. Its compatibility with 384-well formats makes it particularly attractive for drug screening and biomarker validation.

While developed here for alternative splicing analysis, rPCR is applicable to any scenario involving the quantification of relative nucleic acid abundance. Examples include SNP genotyping, mutation frequency estimation in heterogeneous samples, and the detection of viral quasispecies. Clinical applications may involve the detection of low-level drug resistance mutations in infectious diseases or the quantification of isoform switching in response to treatment. As rPCR requires only standard qPCR equipment and existing assays can be adopted, it may be attractive for clinical laboratories, diagnostics, and high-throughput screening platforms.

In conclusion, rPCR combines the accessibility of qPCR with the precision of a mechanistic correction strategy, thereby providing a practical and accurate method for measuring target ratios in a range of molecular biology and clinical applications. The flexibility, speed, and minimal equipment requirements of this method make it well-suited for biomarker validation, therapeutic development, and large-scale screening efforts.

## ACKNOWLEDGEMENTS

We thank Thorsten Fritzius and Tobias Weiss reading the manuscript and for helpful discussions. We are grateful to Eleonora Maino for her help with digital PCR experiments.

## AUTHOR CONTRIBUTIONS

Jochen Kinter: Conceptualization, Data curation, Formal analysis, Investigation, Visualization, Methodology, Project administration, Software, Supervision, Validation, Visualization, Writing—original draft. Adeline Stiefvater: Data curation, Formal analysis, Investigation, Visualization, Methodology. Samira Osterop: Data curation, Formal analysis, Investigation, Investigation, Methodology. Michael Sinnreich: Project administration, Writing—review & editing.

## SUPPLEMENTARY DATA

Supplementary data is available.

## DATA AVAILABILITY

The data underlying this article are available in the article and in its online supplementary material.

## References

1. Kubista, M. et al. The real-time polymerase chain reaction. Molecular Aspects of Medicine 27, 95–125 (2006).

2. Holland, P. M., Abramson, R. D., Watson, R. & Gelfand, D. H. Detection of specific polymerase chain reaction product by utilizing the 5’ 3’ exonuclease activity of Thermus aquaticus DNA polymerase. Proc Natl Acad Sci U S A 88, 7276–7280 (1991).

3. Kuang, J., Yan, X., Genders, A. J., Granata, C. & Bishop, D. J. An overview of technical considerations when using quantitative real-time PCR analysis of gene expression in human exercise research. PLoS One 13, e0196438 (2018).

4. Espy, M. J. et al. Real-time PCR in clinical microbiology: applications for routine laboratory testing. Clin Microbiol Rev 19, 165–256 (2006).

5. Kuypers, J. & Jerome, K. R. Applications of Digital PCR for Clinical Microbiology. J Clin Microbiol 55, 1621–1628 (2017).

6. Men, Y. et al. Digital Polymerase Chain Reaction in an Array of Femtoliter Polydimethylsiloxane Microreactors. Anal. Chem. 84, 4262–4266 (2012).

7. Sykes, P. J. et al. Quantitation of targets for PCR by use of limiting dilution. Biotechniques 13, 444–449 (1992).

8. Mao, X., Liu, C., Tong, H., Chen, Y. & Liu, K. Principles of digital PCR and its applications in current obstetrical and gynecological diseases. Am J Transl Res 11, 7209–7222 (2019).

9. Ruijter, J. M. et al. Evaluation of qPCR curve analysis methods for reliable biomarker discovery: bias, resolution, precision, and implications. Methods 59, 32–46 (2013).

10. Kalle, E., Kubista, M. & Rensing, C. Multi-template polymerase chain reaction. Biomolecular Detection and Quantification 2, 11–29 (2014).

11. Gilliland, G., Perrin, S., Blanchard, K. & Bunn, H. F. Analysis of cytokine mRNA and DNA: detection and quantitation by competitive polymerase chain reaction. Proc Natl Acad Sci U S A 87, 2725–2729 (1990).

12. Siebert, P. D. & Larrick, J. W. Competitive PCR. Nature 359, 557–558 (1992).

13. Becker-André, M. & Hahlbrock, K. Absolute mRNA quantification using the polymerase chain reaction (PCR). A novel approach by a PCR aided transcript titration assay (PATTY). Nucleic Acids Res 17, 9437–9446 (1989).

14. Steffan, R. J. & Atlas, R. M. POLYMERASE CHAIN REACTION: Applications in Environmental Microbiology. Annual Review of Microbiology 45, 137–161 (1991).

15. Surveys of gene families using polymerase chain reaction: PCR selection and PCR drift. Syst Biol. 1994;43:250–261

16. Meiser, L. C. et al. Reading and writing digital data in DNA. Nat Protoc 15, 86–101 (2020).

17. Park, S. H. et al. Comprehensive analysis and accurate quantification of unintended large gene modifications induced by CRISPR-Cas9 gene editing. Science Advances 8, eabo7676 (2022).

18. Polz, M. F. & Cavanaugh, C. M. Bias in Template-to-Product Ratios in Multitemplate PCR. Applied and Environmental Microbiology 64, 3724–3730 (1998).

19. Alvarez, M. J., Depino, A. M., Podhajcer, O. L. & Pitossi, F. J. Bias in Estimations of DNA Content by Competitive Polymerase Chain Reaction. Analytical Biochemistry 287, 87–94 (2000).

20. Y. Trick, A., et al. High resolution estimates of relative gene abundance with quantitative ratiometric regression PCR (qRR-PCR). Analyst 146, 6463–6469 (2021).

21. Tomita, K., Indo, H. P., Sato, T., Tangpong, J. & Majima, H. J. Development of a sensitive double TaqMan Probe-based qPCR Angle-Degree method to detect mutation frequencies. Mitochondrion 70, 1–7 (2023).

22. Tajima, H. et al. The development of novel quantification assay for mitochondrial DNA heteroplasmy aimed at preimplantation genetic diagnosis of Leigh encephalopathy. J Assist Reprod Genet 24, 227–232 (2007).

23. Herrendorff, R. et al. Identification of Plant-derived Alkaloids with Therapeutic Potential for Myotonic Dystrophy Type I. J Biol Chem 291, 17165–17177 (2016).

24. Foff, E. P. & Mahadevan, M. S. Therapeutics development in myotonic dystrophy type 1. Muscle Nerve 44, 160–169 (2011).

25. Nakamori, M. et al. Splicing biomarkers of disease severity in myotonic dystrophy. Ann Neurol 74, 862–872 (2013).

26. Wojciechowska, M. et al. Quantitative Methods to Monitor RNA Biomarkers in Myotonic Dystrophy. Sci Rep 8, 5885 (2018).

27. Jenquin, J. R. et al. Molecular characterization of myotonic dystrophy fibroblast cell lines for use in small molecule screening. iScience 25, 104198 (2022).

28. Fritzson, P. et al. The OpenModelica integrated environment for modeling, simulation, and model-based development. 10.4173/MIC.2020.4.1 (2022) doi:10.4173/MIC.2020.4.1.

29. Asghar, A. The OpenModelica Integrated Modeling, Simulation, and Optimization Environment. Proceedings of The American Modelica Conference 2018, October 9-10, Somberg Conference Center, Cambridge MA, USA 10.3384/ECP18154206 (2022) doi:10.3384/ECP18154206.

30. Fritzson, P., Ulfhielm, E., Belic, A., Fransson, M. & Green, H. Biochemical mathematical modeling with modelica and the biochem library. in Proceedings of 6th International Conference on Applied Mathematics (APLIMAT 2007) 147–159 (2007).

31. Arandel, L. et al. Immortalized human myotonic dystrophy muscle cell lines to assess therapeutic compounds. Dis Model Mech 10, 487–497 (2017).

32. Schölzel, C., Blesius, V., Ernst, G. & Dominik, A. Characteristics of mathematical modeling languages that facilitate model reuse in systems biology: a software engineering perspective. npj Syst Biol Appl 7, 27 (2021).

33. Mehra, S. & Hu, W.-S. A kinetic model of quantitative real-time polymerase chain reaction. Biotechnology and Bioengineering 91, 848–860 (2005).

34. Gevertz, J. L., Dunn, S. M. & Roth, C. M. Mathematical model of real-time PCR kinetics. Biotechnology and Bioengineering 92, 346–355 (2005).

35. Blomquist, T. M. et al. Targeted RNA-Sequencing with Competitive Multiplex-PCR Amplicon Libraries. PLOS ONE 8, e79120 (2013).

36. Blomquist, T. M. et al. Targeted RNA-Sequencing with Competitive Multiplex-PCR Amplicon Libraries. PLOS ONE 8, e79120 (2013).

37. Harvey, S. E., Lyu, J. & Cheng, C. Methods for characterization of alternative RNA splicing. Methods Mol Biol 2372, 209–222 (2021).

38. Wojciechowska, M. et al. Quantitative Methods to Monitor RNA Biomarkers in Myotonic Dystrophy. Sci Rep 8, 5885 (2018).

39. Sun, B., Tao, L. & Zheng, Y.-L. Simultaneous Quantification of Alternatively Spliced Transcripts in a Single Droplet Digital PCR Reaction. BioTechniques 56, 319–325 (2014).

40. Alamancos, G. P., Agirre, E. & Eyras, E. Methods to study splicing from high-throughput RNA sequencing data. Methods Mol Biol 1126, 357–397 (2014).

41. Draper, B. J., Dunning, M. J. & James, D. C. Selecting differential splicing methods: Practical considerations for short-read RNA sequencing. F1000Res 14, 47 (2025).

