## Supplemental Figures for "ratioPCR delivers precise real-time quantification from multi-template PCR via mechanistic bias correction"

**Supplementary Figure 1**

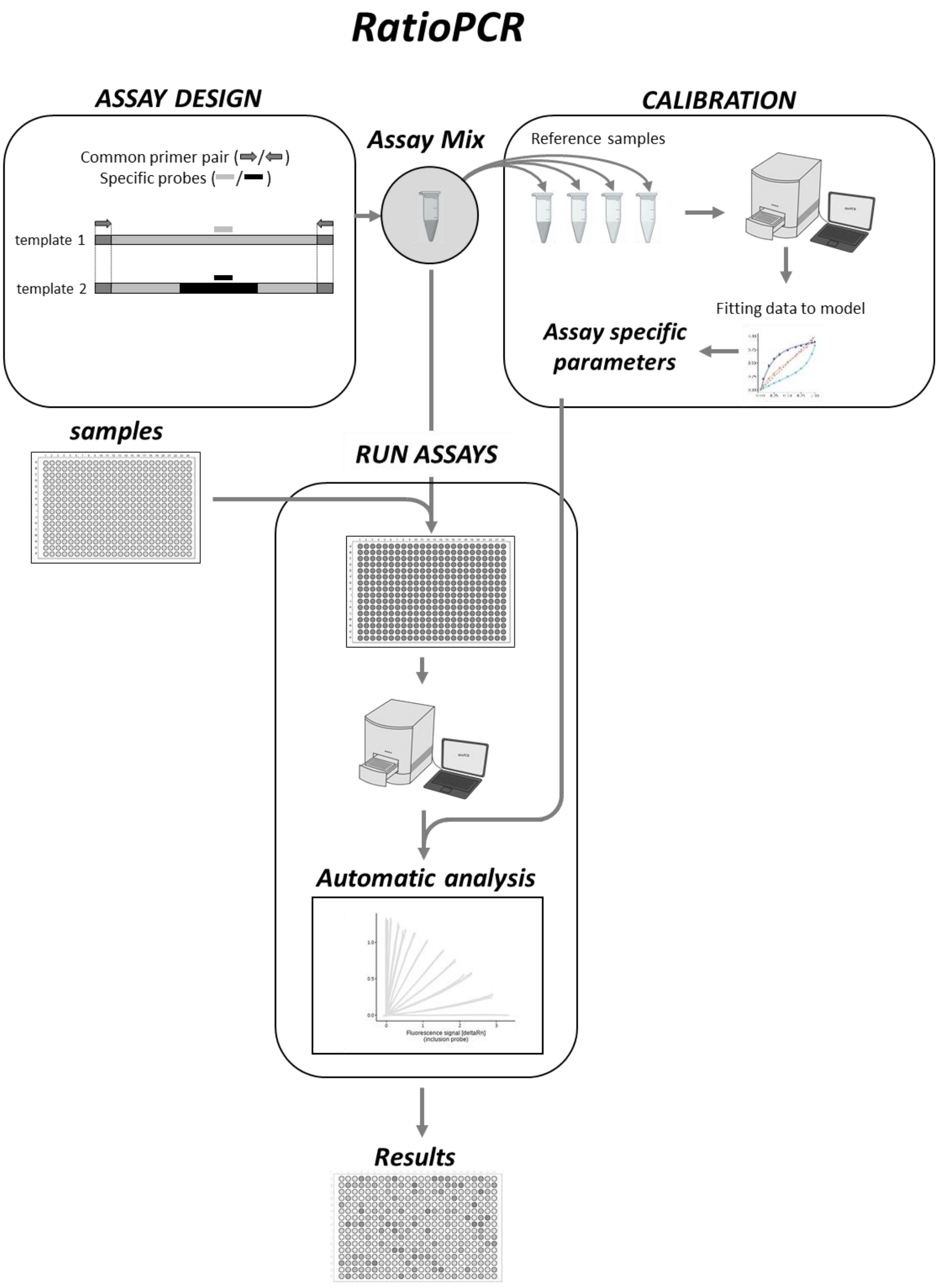


**Supplemental Figure 1: Workflow of establishing and running an rPCR assay.**
The requirement of a rPCR is a common primer pair for both templates and template specific probes. Primer and probe design tools developed for qPCR can be used to design the assay. In addition to the primer and probes synthetic templates should be ordered. Using defined rations of the synthetic templates the assay specific parameters are calibrated. All the rPCR experiments can be performed with any standard qPCR machine. After running rPCR the obtained ratio data are corrected using the parameter obtained by the calibration experiment.

#### Supplementary Figure 2

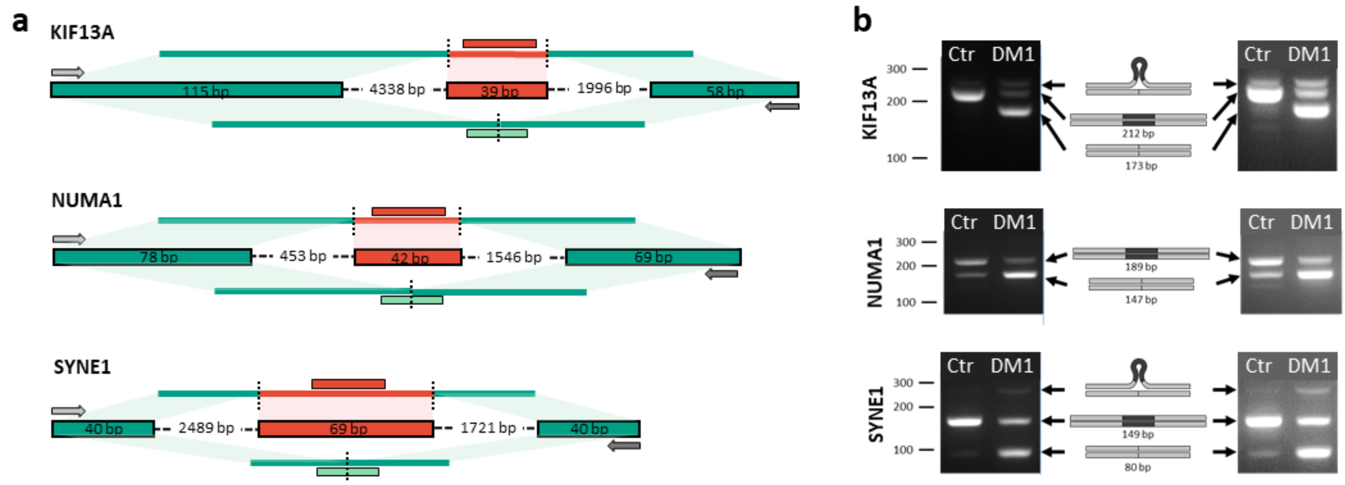


**Supplemental Figure 2: Primer design and validation.**
**a**, Genomic context and assay design for three alternatively spliced genes (KIF13A, NUMA1, SYNE1). Red boxes mark the alternative exons. Primer binding sites (arrows) and probe locations (red: inclusion; green: exclusion) are shown. **b**, Agarose gel electrophoresis of RT–PCR products from control and DM1 myoblast RNA reveals altered splicing patterns. Heteroduplexes between isoforms form slower-migrating bands.

**Supplementary Figure 3**

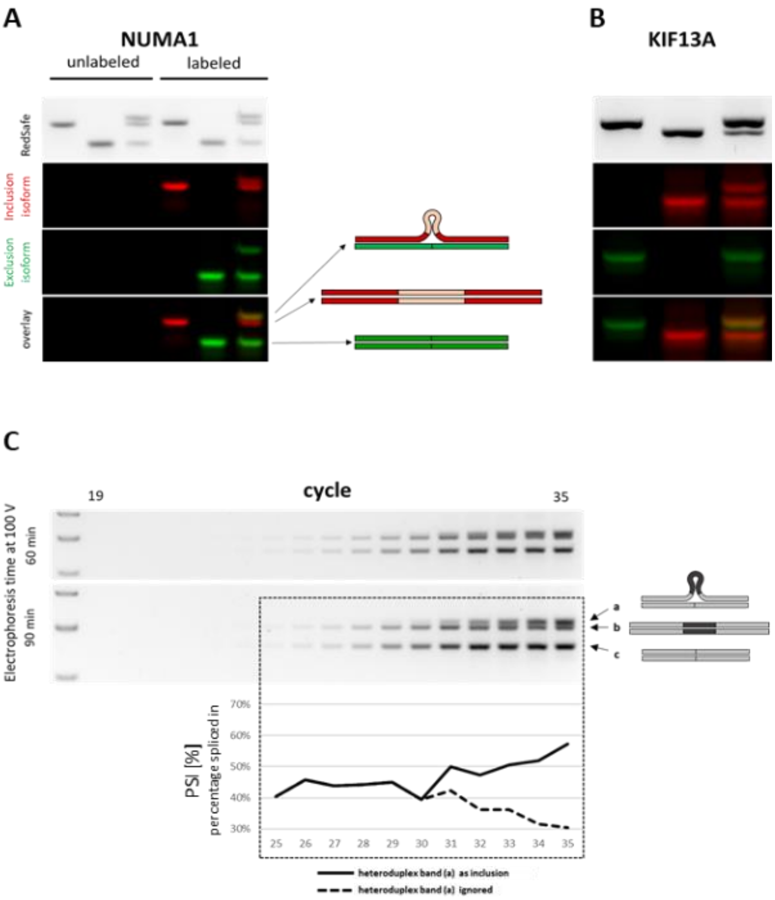


**Supplemental Figure 3: Heteroduplex formation of inclusion and exclusion isoforms.**
A) Single isoforms were amplified using fluorescence labeled primer. Re-annealing experiments were performed either with pure isoform samples or a 1:1 mixture of exclusion and inclusion isoforms. Samples were heated at 95 °C for 15 seconds and re-annealed for 45 seconds at 60°C. After re-annealing samples were analyzed by agarose gel-electrophorese and different isoforms were detected by the distinct fluorescent dyes. For NUMA1 heteroduplex formation can already be visualized by using a general DNA dye like RedSsafe. The heteroduplex are visible ad an additional slower migrating band. B) Identical assay as A) using KIF13A templates and primers. No additional heteroduplex band can be visualized using the general DNA dye RedSafe. The electrophoretic conditions (agarose content, time, etc) are not able to separate the heteroduplex population from inclusion isoform. The formation of heteroduplex is nevertheless confirmed by the shift of exclusion strands ( green) to the mixed inclusion & heteroduplex band.

**Supplementary Figure 4**

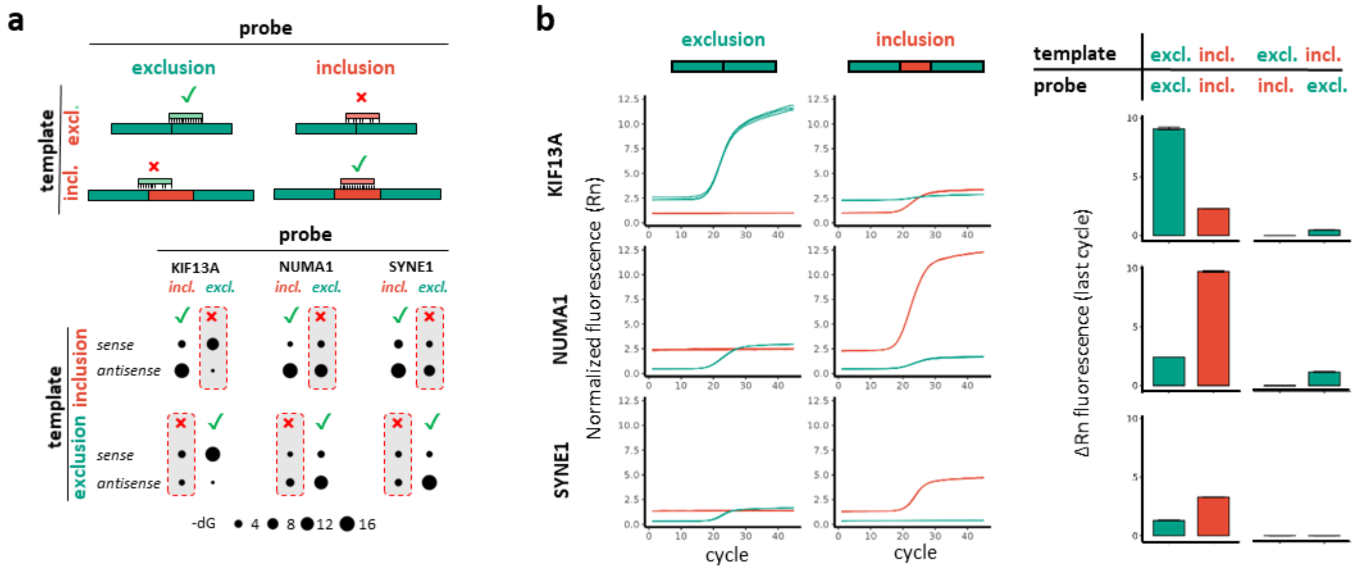


**Supplemental Figure 4: Analysis of cross-reactivity of isoform specific probes.**
**f**, Predicted Gibbs free energy (ΔG) of probe–template binding shows high specificity for inclusion probes and variable cross-reactivity for exclusion probes. **g**, Experimental validation of probe specificity using synthetic isoform templates. Bar graphs show endpoint fluorescence (ΔRn). Inclusion probes exhibit no cross-reactivity; exclusion probes display target-dependent off-target binding, most notably for NUMA1.

### Supplementary Figure 5

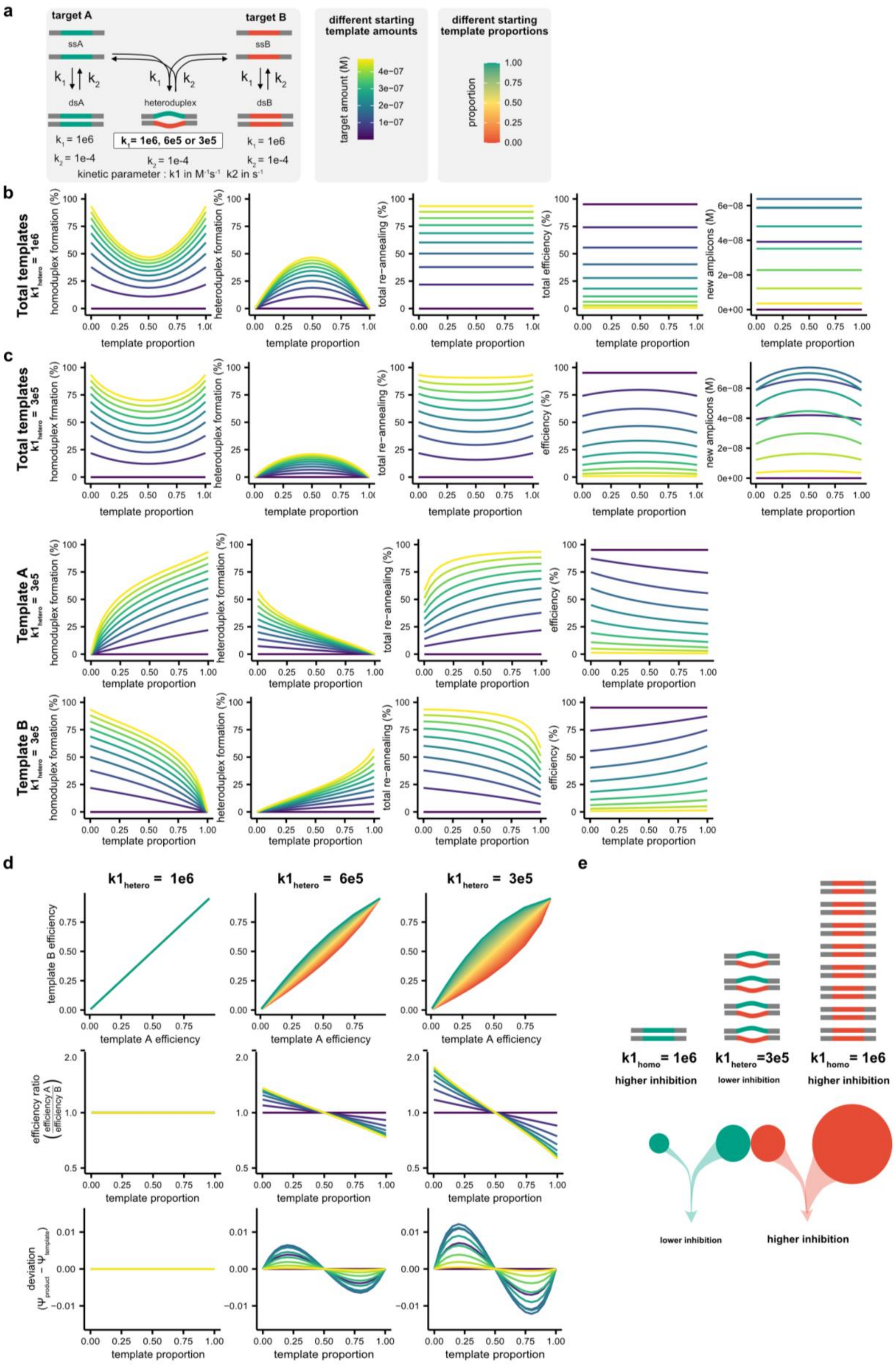

**Supplemental Figure 5: Simulation of equimolar drift due to different annealing properties of heteroduplexes.**

**a**, Single cycles were simulated using different total template amounts, different heteroduplex formation kinetics, and different template proportions (ratios). The simulations was performed with an association rate for the homoduplexes of 1e6 $M^{-1}s^{-1}$ and different association rates for heteroduplexes (1e6, 6e5, and 3e5 $M^{-1}s^{-1}$). Different starting template amount and starting proportion are visualized with two different color codes. **b**, Simulation for equal association rates for homo- and heteroduplexes formation by re-annealing. The fraction of homoduplex and heteroduplex formation by re-anealing depends on the template proportions. At equal template amounts, the formation of homoduplexes is minimal and the formation of heteroduplex is maximal. The fraction of all re-annealing duplex is increasing with increasing template amounts. The amplification efficiency in a cycle is decreasing with increasing template concentrations. With equal association rates, total re-annealing and amplification efficiency is not dependent on template proportions. **c**, A more realistic simulation is that heterodplex have lower affinity than homoduplexes due to mismatches. Therefore, we run the simulation with heteroduplex association rates lower than rates of the homoduplexes. Again, the fraction of homoduplex and heteroduplex formation by re-annealing depends on the template proportions. At equal template amounts, the formation of homoduplexes is minimal and the formation of heteroduplex is maximal. The total re-annealing fractions and the total amplification is dependent on the template ratio. The efficiency is highest at 1:1 template ratios. With increasing relative amounts of the single templates A and template B, the fraction of re-annealed molecules in homoduplexes increases and the fractions of molecules in heteroduplexes decreases. In addition, the fraction of molecules participating in re-annealing increase and the specific template efficiency decreases with increasing relative template amounts. The specific efficiency of both templates is equal at equal template amounts. **d**, At equal re-annealing rates for homo- and heteroduplexes the efficiency for both templates is equal independent of the template proportions. Decreasing the association rate of heteroduplexes leads to proportion dependent difference in amplification efficiency of each template, with the exception of equal template amounts (1:1). These differences in efficiency leads to a proportion dependent template-to-product bias resulting in a drift towards equimolar amounts. **e**, In a multi-template PCR effiency is enhanced by the formation of heteroduplexes. The fraction of molecules in heteroduplexes is higher for the less abundant template, leading to higher efficiency and a drift to equimolar amounts.
