## Supplemental Notes for "ratioPCR delivers precise real-time quantification from multi-template PCR via mechanistic bias correction"

### Supplementary Note 1: Kinetic Simulation Model of ratioPCR Amplification Dynamics

To better understand the amplification behavior and ratio distortion observed in ratioPCR (rPCR), we developed a detailed kinetic simulation model of the coupled amplification process. The model was implemented in Modelica using the OpenModelica platform and the BioChem library. It simulates the full reaction kinetics of two DNA targets amplified using common primers and detected using target-specific probes.

#### Model Overview

The model describes the kinetics of the following mechanisms:

- Primer hybridization to each template strand (target A and B)
- Extension of primers by DNA polymerase to form double-stranded amplicons
- Denaturation and reannealing of single-stranded templates
- Heteroduplex formation
- Competition for primers among targets during amplification
- Binding of fluorescent probes to specific amplicons
- Cross-hybridization of the probes to “wrong” templates

A total of 781 differential and algebraic equations were used to simulate these reactions, of which 495 are algebraically trivial (e.g., mass balances, conservation constraints). Kinetic parameters were adapted from published literature on PCR enzyme kinetics, oligonucleotide binding affinities, and strand displacement rates.

#### Key Features and Assumptions

- Both targets share a common primer pair.
- Fluorescent probe binding occurs independently from polymerase extension.
- The efficiency of amplification is governed by template rehybridization and primer availability
- Sequence-specific rehybridization rates affect amplification efficiency and contribute to target-specific bias.
- Amplification bias increases with template.

#### Simulation Conditions

- Initial template concentrations were set to represent various known ratios (e.g., 0.1 to 0.9).
- Simulations were run for 40 cycles using a standard thermal cycling profile.
- Primer concentrations, enzyme kinetics, and probe affinities were adjusted based on typical qPCR reaction conditions.

#### Results

The kinetic simulations recapitulated several experimentally observed phenomena:

- Progressive distortion of the target ratio with increased amplification.
- Amplification slowdown for abundant templates, allowing lower abundance templates to relatively "catch up".
- Nonlinear fluorescence ratio trajectories, consistent with rPCR experimental data.

The results confirm that even with shared primers, sequence-dependent differences in amplification efficiency emerge due to differential reannealing and primer competition. This supports the need for bias correction in rPCR.

#### Motivation for Analytical Model Development

While the kinetic model provides deep mechanistic insight, its complexity makes it impractical for routine use in data analysis and assay calibration. Therefore, we developed a simplified analytical model described in Supplementary Note 2. This model captures key amplification and probe-related biases using a small number of interpretable parameters, enabling efficient curve fitting and correction of measured fluorescence ratios.

#### Software and Availability

The full Modelica code for the kinetic model is available upon request and will be deposited in a public repository upon publication.

Simulation software: OpenModelica 4.0 Library and BioChem Library 2.0.

R was used to visulaize the results.

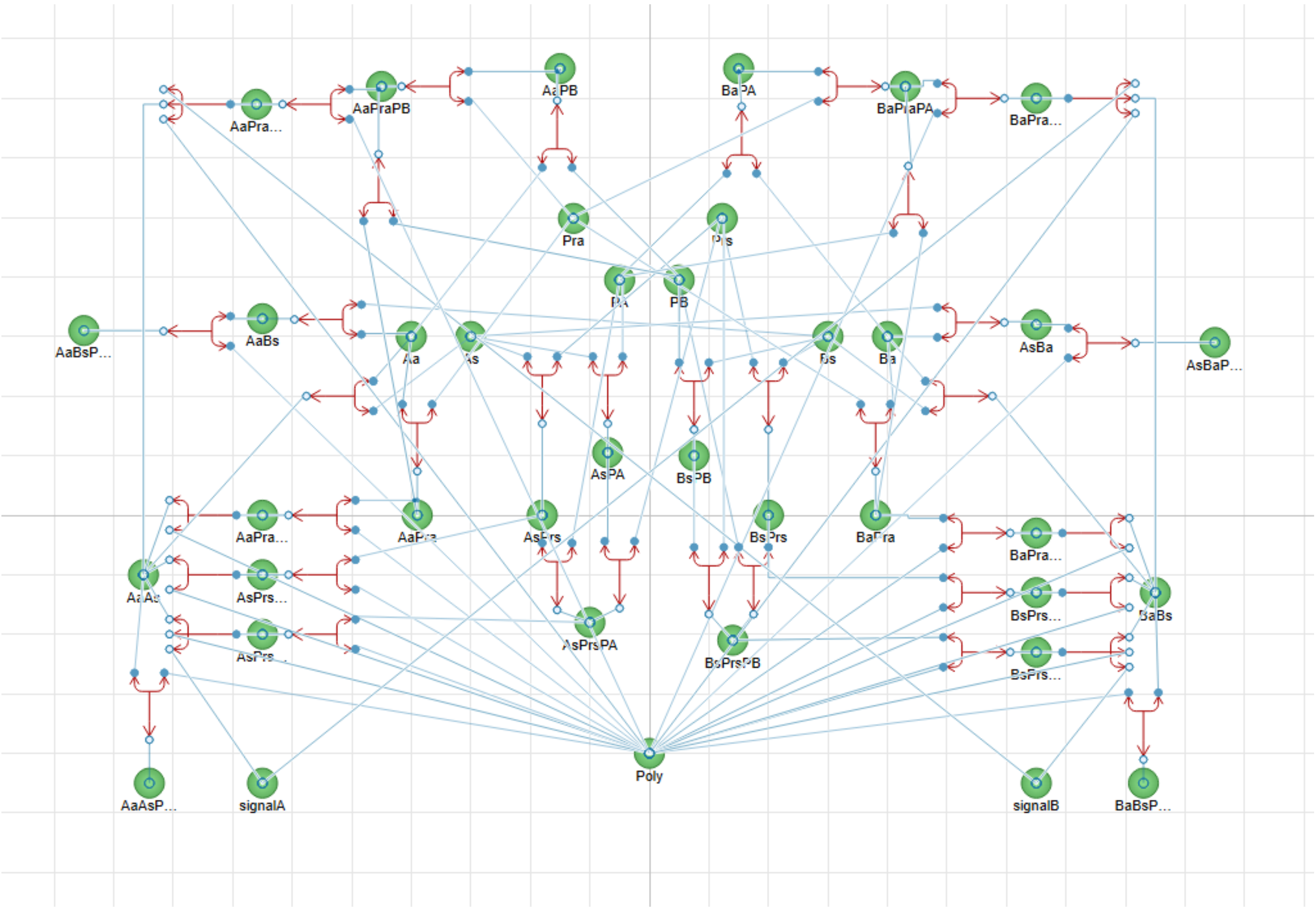


**Supplemental Note 1 Figure A:**
Graphical View (OMEdit) of the kinetic model demonstrating the complexity. Each Green circle represents a substance (either a single molecule or complex). The red arrows represent a reversible or irreversible reaction. The blue lines connect the different substances as reaction partner to a reaction.

**Supplementary Note 2: Mathematical Model and Calibration of ratioPCR Bias**

### Overview

ratioPCR (rPCR) quantifies the relative abundance of nucleic acid targets by comparing fluorescence signals from isoform-specific probes during PCR amplification using shared primers. However, differences in probe intensity, cross-reactivity, and amplification efficiency can bias the measured ratios. Here, we present a theoretical model to correct these biases and describe how to calibrate the model using reference samples.

### Definitions

| Symbol | Description |
|---|---|
| $N_A$ | Number of target A molecules |
| $N_B$ | Number of target B molecules |
| $N_T$ | Number of all molecules |
| $ϙ$ | ratio of two targets (eg. $N_A/N_B$) |
| $ϙ_a$ | apparent $ϙ$ (the biased uncorrected ratio) |
| $\psi$ | proportion of target of interest |
| $\psi_a$ | apparent $\psi$ (the biased uncorrected proportion) |
| $F_A$ | Specific fluorscence signal generated by probe detecting target A |
| $F_B$ | Specific fluorscence signal generated by probe detecting target B |
| $F_T$ | Fluorscence signal generated by a probe or dye detecting all targets |
| $\alpha$ | parameter describing the fluorescence intensity ratio between two probes |
| $r_i$ | The proportion of off-targets recognized by probe A |
| $r_e$ | The proportion of off-targets recognized by probe B |
| $\gamma$ | A term specifying the degree of ratio-dependent efficiency bias. |

### Basic Proportionality Model

In a two-target amplification system:

Ratio:
$$ϙ = \frac{N_A}{N_B} \qquad \text{Eq. 1}$$

Proportion:
$$\psi = \frac{N_A}{N_A + N_B} = \frac{N_A}{N_T} \qquad \text{Eq. 2}$$

Relationship:
$$\psi = \frac{ϙ}{1 + ϙ}\ ;\ ϙ = \frac{\psi}{1 - \psi} \qquad \text{Eq. 3}$$

Under the assumption that fluorescence is proportional to the number of amplicons ($F \propto N$), we can define the apparent quantities as:

$$\text{Ϙ}_a = \frac{F_A}{F_B} \qquad \text{Eq. 4}$$

$$\psi_a = \frac{F_A}{F_A + F_B} = \frac{F_A}{F_T} \qquad \text{Eq. 5}$$

### Sources of Bias

a) Probe-Specific Fluorescence Intensity

Probes differ in signal yield due to fluorophores, quenchers, sequence, and qPCR machine optics. We define $= \frac{k_A}{k_B}$, the relative fluorescence scaling factor between probes.

b) Cross-Reactivity

Probes may bind off-target templates. We model this with coefficients $r_i$ and $r_e$, such that:

- $F_A = k_A(N_A + r_i N_B)$
- $F_B = k_B(N_B + r_e N_A)$

This leads to:

$$\text{Ϙ}_a = \frac{F_A}{F_B} = \alpha \frac{\text{Ϙ} + r_i}{1 + r_e \text{Ϙ}} \qquad \text{Eq. 6}$$

$$\psi_a = \frac{\alpha\psi(1 - r_i) + \alpha r_i}{\psi(\alpha - \alpha r_i - 1 + r_e) + \alpha r_i + 1} \qquad \text{Eq. 7}$$

c) Target-Specific Amplification Efficiency

Differences in sequence and re-hybridization rates cause target-specific amplification biases. If target A has amplification efficiency $E_A$ and target B has $E_B$, we define $m = E_A / E_B$.

The apparent proportion becomes:

$$\psi_a = m_C \psi \qquad \text{Eq. 8}$$

d) Ratio-Dependent Efficiency Bias

As inhibitory reannealing is more pronounced for more abundant templates, they become less efficiently amplified. We introduce a phenomenological correction factor:

$$z = \gamma^{1-2\psi} \qquad \text{Eq. 9}$$

This term models the efficiency convergence toward a 1:1 ratio during late PCR cycles.

**Final Corrected Model**

Combining probe intensity, cross-reactivity, and efficiency bias, we derive a general correction equation:

$$\psi_a = \frac{\psi\alpha\gamma^{(1-2\psi)} - \psi r_i + r_i}{\psi\,\alpha\gamma^{(1-2\psi)}(1 + r_e) - \psi r_i - \psi + r_i + 1} \qquad \text{Eq. 10}$$

**Calibration Procedure**

Reference samples with known target ratios ($\psi$) are amplified by rPCR, and the resulting $\psi_a$ values are fitted to the analytical model using nonlinear regression (e.g., Levenberg-Marquardt algorithm). Once calibrated, assay-specific parameters $\alpha$, $r_i$, $r_e$, $\gamma$ can be applied to correct experimental data.

An open-source R package was developed to automate this calibration and correction process.

**Conclusion**

This model offers a simplified mechanistic, flexible, and calibratable approach for correcting bias in ratioPCR measurements. Its implementation enables accurate relative quantification from standard qPCR instruments, enhancing assay performance across diagnostic and research applications.
